# Analysis of Independent Differences (AID) detects complex thermal proteome profiles independent of shape and identifies candidate panobinostat targets

**DOI:** 10.1101/751818

**Authors:** Alexandra Panov, Steven P. Gygi

## Abstract

Identifying global cellular targets of small molecules is a challenge for drug discovery. Thermal proteome profiling (TPP) is a recent technique that uses quantitative proteomics to identify all small molecule protein targets in a single experiment. One current TPP analysis method relies on two major assumptions: sigmoidal melting curve behavior and that intra-condition dependencies preclude an independent and identically distributed model. Herein, we use a previously published panobinostat TPP dataset to show that these assumptions do not hold true and present a novel, shape-independent method, named Analysis of Independent Differences (AID). For each temperature, AID models the differences between conditions of fractions of non-denatured protein as an independent Normal distribution, resulting in a Multivariate Normal observation for each protein. The log of a Multivariate Normal *p*-value ranks the proteins from most to least likely shifted, and individual Normal *p*-values within each protein allow for qualitative inspection. Applying AID to the panobinostat dataset revealed known targets in the top 3% of most shifted proteins, as well as candidate targets involved in myeloid leukocyte activation. AID detects complex melting profiles and can be extended to any number of temperature channels, ligand-protein or protein-protein interactions, or general curve data for deeper biological insight.

## Introduction

Identifying physiologically relevant direct and indirect protein targets of small molecules remains a great challenge for modern day drug discovery. Chemical probes are often used to validate protein targets, but concerns have surfaced regarding probe quality, sufficient controls, and subsequent experimental conclusions^1,2,3^. To address this challenge, thermal proteome profiling (TPP), an extension of the cellular thermal shift assay (CETSA), has been successfully employed to monitor small molecule interactions within whole cells and cell lysate^4,5,6^. TPP categorizes small molecule interactors by overall stabilization or destabilization of a given protein, which is observed as a shift in the protein’s melting curve. Whole cells or lysate are treated with a small molecule of interest, which along with controls, are subjected to a temperature gradient and examined for solubility. Multiplexed quantitative mass spectrometry techniques are used to generate thousands of protein melting curves in a single experiment. TPP has been further applied to probe interactions between endogenous ligands and proteins^7^, protein-protein interactions^8^, and overall proteome stability in prokaryotes and eukaryotes^9,10^.

The most common method for analyzing TPP data has both a parametric and non-parametric approach^4,6,11^. The parametric approach fits a sigmoidal function to the data and assesses if there is a difference between the melting temperature, T_m_, of control and treatment conditions, where T_m_ is the temperature at which the fraction of non-denatured protein equals 0.5. The differences between T_m_ are binned by increasing slope of the fitted curve, and a *z*-test is performed within each bin to assign a *p*-value to the difference in T_m_ of each protein. Qualitative filters applied throughout the process result in a rigor-variance tradeoff; while lenient filters allow more curves to be analyzed, increasing the number of curves with a poor sigmoidal fit likewise increases the variance, which decreases confidence during hypothesis testing^12^. Moreover, if T_m_ is strictly limited to the temperature at which the fraction of non-denatured protein equals 0.5, then a shift in T_m_ may not accurately represent a change in protein thermostability, or T_m_ may not be calculable from the data. For these reasons, Childs et al. (2018) developed a non-parametric analysis of response curves (NPARC)^11^. NPARC compares the sum of squared residuals of a null model against that of an alternative model, which measures a difference in overall melting curve shape via an F-statistic. The mean function chosen for the null and alternative models is a sigmoidal function derived from chemical denaturation theory^4^. Both approaches assume a large majority of protein melting curves follow an approximately sigmoidal shape, though deviations from this behavior are common^11,13^. NPARC further assumes that intra-condition dependencies preclude an independent and identically distributed model. If either of these assumptions do not hold true, the statistical power of the analysis will be compromised.

Two additional TPP analysis methods have been informally proposed, called Proteome Integral Stability Alteration (PISA)^13^ and TargetSeeker-MS^14^. PISA collapses melting curve data into a single statistic by physically mixing samples in each condition after exposure to a temperature gradient, which are then analyzed by mass spectrometry. A *t*-test is performed to detect a significant difference in protein abundance between conditions, representing a shift in protein thermostability. The experimental design allows for five replicates in a single TMT 10-plex. However, this method also relies on the assumption of sigmoidal melting curve behavior. In addition, the subsequent single data point for a given protein is blind to qualitative inspection, which may increase the Type I or Type II error rate or decrease power. TargetSeeker-MS employs a Bayesian inference-based approach to compare the probability that a fractionation profile of a given protein in the control condition is different from that in the treatment condition. Fractionation profiles are compared by subtracting scaled Euclidean distances from 1, yielding two similarity values, one for each condition. Because a Bayesian approach warrants constructing a reasonable prior distribution, TargetSeeker-MS requires at least 4 control replicates and 2 treatment replicates, requiring 5 full days of instrument time. While this method does not assume sigmoidal melting curve behavior, it is impractically resource-intensive.

Herein, we demonstrate limitations of previous TPP analysis methods and present a novel method capable of parametrically analyzing all melting curves regardless of shape, named Analysis of Independent Differences (AID). This method estimates the underlying null distribution by circumventing dependencies within each condition, and is more sensitive for detecting subtle and complex melting curve shifts. To demonstrate the method, we analyze a previously published gold standard TPP dataset that investigates panobinostat, a therapeutic for multiple myeloma^6^. Not only does the presented method identify previously known targets, including HDACs, it also identifies many other proteins, including those involved in leukocyte activation, providing further biological insight into the therapeutic efficacy of panobinostat.

## Methods

The dataset used is from Franken et al. (2015) and is publicly available. Two replicates of thermal proteome profiling of panobinostat were performed in intact K562 cells, a human chronic myelogenous leukemia cell line, as described^6^. Briefly, TMT-labelled LC-MS/MS spectra were analyzed using *isobarQuant*, a software package provided by Franken et al. (2015), and Mascot, a common search engine. Peptide fold changes are calculated from corrected reporter ion intensities, and subsequent protein fold changes are calculated using a sum-based bootstrapping approach. The protein fold changes are then normalized to a value of 1 in the lowest temperature channel of each protein, such that most normalized fold changes fall between 0 and 1.

Examining both replicates of the panobinostat data showed that only approximately 42% of proteins exhibit a maximum protein abundance value in the lowest temperature channel. Approximately 25% of proteins exhibit a maximum abundance value in the second lowest temperature channel, which is not surprising, as many proteins are stable at temperatures near 40°C. For these reasons, we instead normalized to a value of 1 in the maximum protein abundance temperature channel for each protein and did not employ further normalization.

Only one replicate was necessary for AID analysis. We proceeded with the second replicate due to a larger number of proteins shared between control and treatment conditions (~300 more proteins).

Sigmoidal curve fitting was performed using a function derived from chemical denaturation theory^4^:

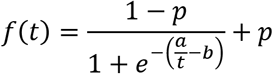

where *t* is temperature (°C), *p* is the plateau of the curve, and *a* and *b* are constants. The value of *p* was limited to the interval [0,1], in accordance with normalization.

Data analysis was performed in R using the *mvtnorm* package. The analysis was written in R and as a Shiny App. The R script is open-source and available at https://github.com/alex-bio/TPP, and the Shiny App is available at https://gygi.med.harvard.edu/software.

## Results and Discussion

We assessed the validity of the two assumptions made by the most common TPP analysis method, namely sigmoidal melting curve behavior and that intra-condition dependencies preclude an independent and identically distributed model, using a previously published gold standard TPP dataset^6^.

First, because both approaches of the method assume that most protein melting curves are sigmoidal in shape, curves from a simple random sample were fitted to a sigmoidal function derived from chemical denaturation theory^4^, as shown in Figure 1. The coefficient of determination, R^2^, values from the fit demonstrate that while some curves are approximately sigmoidal in shape, complex curves are non-trivially present.

**Figure 1.**
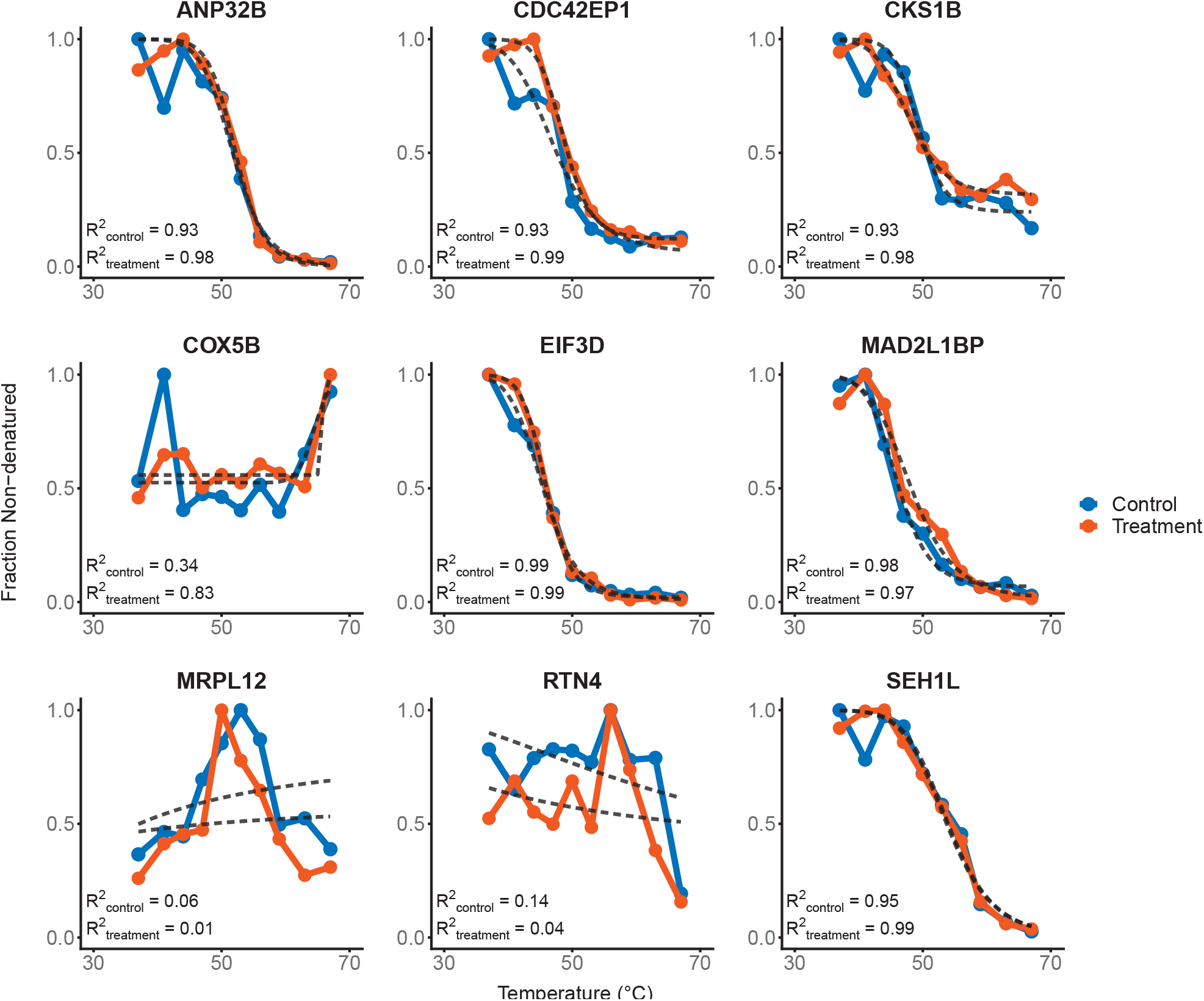
A random sample of melting profiles. Both control and treatment curves for each protein were fitted to a sigmoidal function derived from chemical denaturation theory^4^, shown as dotted gray lines. The R^2^ values are shown, ranging from 0.01 to 0.99.

Second, dependencies within the data were examined. In particular, intra-condition dependencies between the fractions of non-denatured protein add a substantial layer of complexity, as illustrated by the correlation matrices in Figure 2A. Because the global normalization scheme from Savitski et al. (2014) is frequently employed before analyzing TPP datasets, the dependencies of data after global normalization were also examined (Figure 2B). This normalization method significantly alters inherent data dependencies, which would have unintended effects when choosing a null distribution, and thus was not used further. Circumventing dependencies is possible without alteration by examining the *differences* between the control and treatment fractions of non-denatured protein for each temperature (Figure 2C). Specifically, the middle 95% of differences were examined further because of a few extreme outliers, as discussed below. In this case, uncorrelated data strongly suggest an underlying null distribution exists and may have calculable parameters.

**Figure 2.**
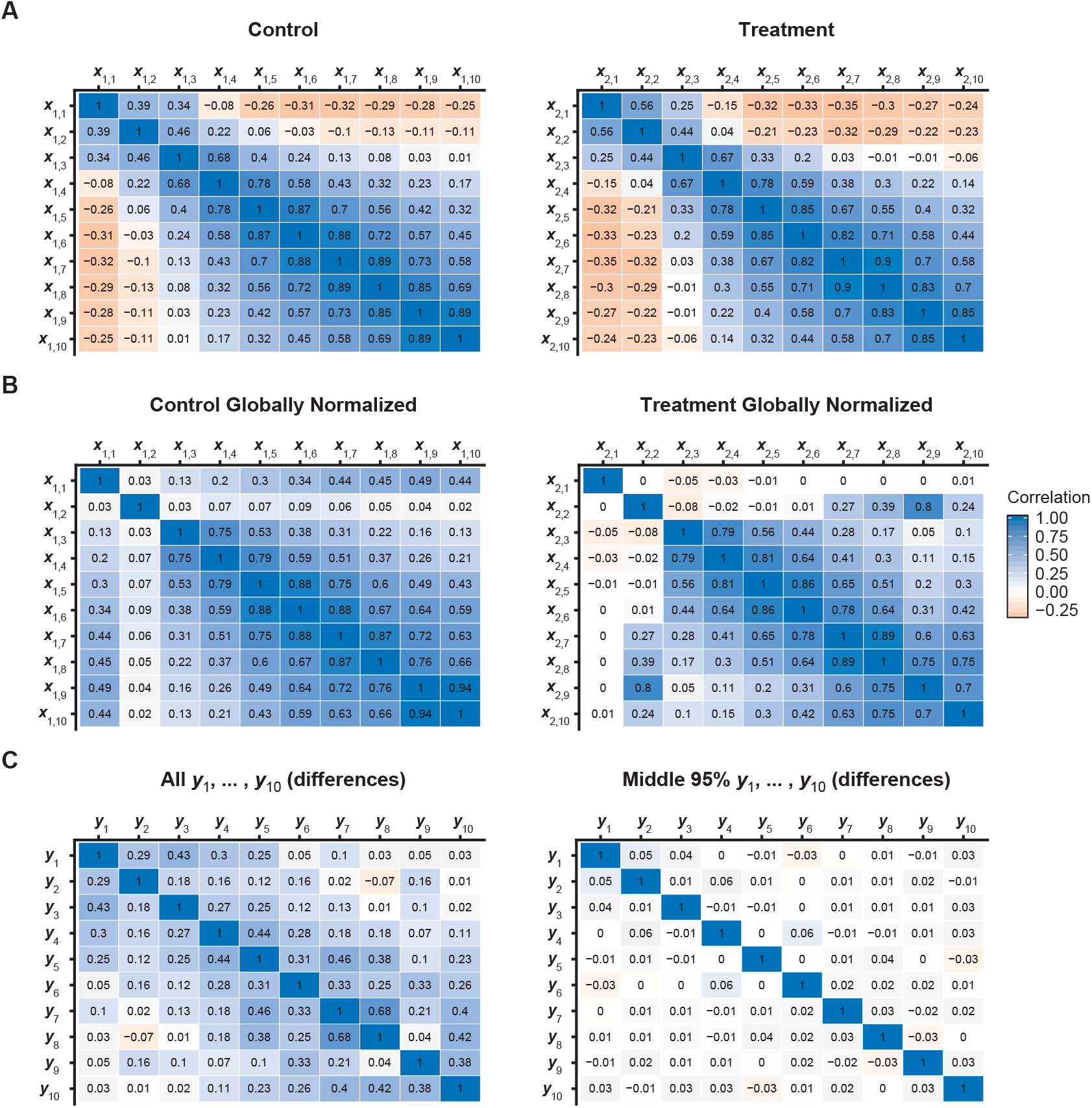
Correlation matrices. (A) Correlation matrix of the fractions of non-denatured protein, or *x_i,j,k_*s, for protein *i* across each temperature *k* for the control condition, *j* = 1 (left), and the treatment condition, *j* = 2 (right), of the panobinostat TPP dataset^6^. (B) Correlation matrix as in (A), except after applying the global normalization scheme from Savitski et al. (2014). (C) Correlation matrix of all the differences between conditions of fractions of non-denatured protein, or *y_i,j_*s, where *y_i,j_* = *x*_*i*,treatment,*k*_ − *x*_*i*,control*,k*_, across each temperature (left). The middle 95% of differences between conditions of fractions of non-denatured protein, or *y_i,j_*s, across each temperature (right).

As a result of both assumptions breaking down, we asked if an alternative statistical test could be used. The lack of dependencies from Figure 2C indicates that the differences between conditions of fractions of non-denatured protein for each temperature may be treated as independent trials from a similar distribution. The fractions of non-denatured protein are represented here as *x_i,j,k_*, for the fraction non-denatured of protein *i*, condition *j*, and temperature *k*. As an example, *x*_10,1,3_ would indicate the fraction of non-denatured protein of the tenth protein in the first condition in the third temperature channel. A histogram of the differences between conditions, *y_i,j_*, for protein *i* and temperature *j*, where *y_i,j_* = *x*_*i*,treatment,*k*_ − *x*_*i*,control,*k*_, was plotted in Figure 3A (blue). As an artifact of normalization, there is a slightly higher peak at exactly zero due to differences between the maximum channels of both conditions, which is not shown. This peak does not affect any further calculations.

**Figure 3.**
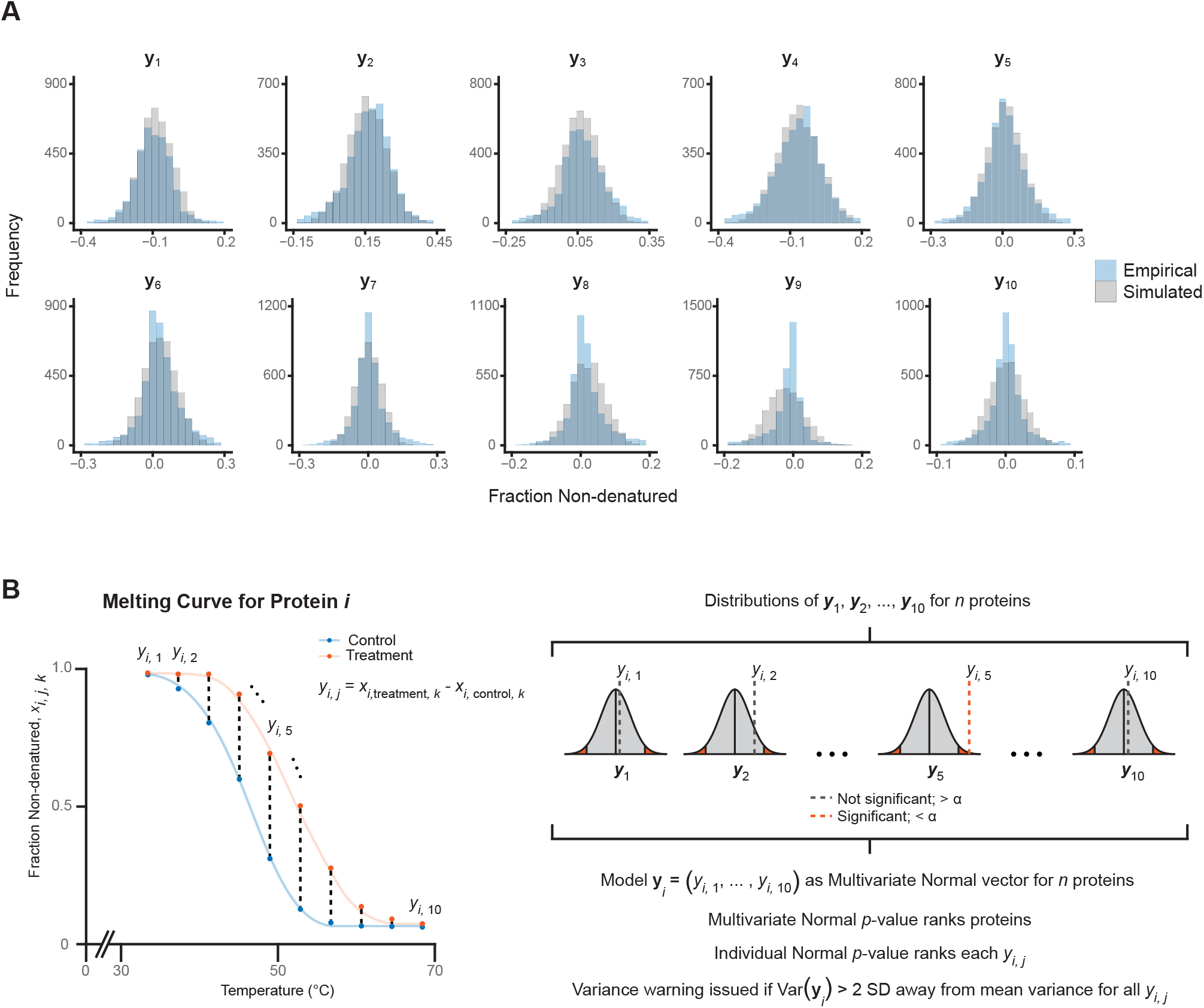
(A) Histograms illustrating differences between conditions of fractions of non-denatured protein, or *y_i,j_*s, across each temperature. The empirical data from the panobinostat TPP dataset^6^ is shown in blue, and simulated data from a fitted Normal distribution is in gray. (B) Diagram of Multivariate Normal modeling. For each protein, the difference between conditions of fractions of non-denatured protein, *y_i,j_*s (black dotted lines), are calculated for each temperature. A vector consisting of all *y_i,j_*s for a given protein is modeled as a Multivariate Normal observation, where each *y_i,j_* is Normally distributed among *y_i,j_*s from all other proteins. Multivariate Normal *p*-values are assigned to each protein, and the log of the *p*-value rank orders the proteins from most likely to least likely shifted.

For each temperature, the differences across all proteins, all *y_i,j_*s, appeared Normally distributed. The law of large numbers grants confidence that approximating a distribution is feasible because of such a large sample size (*n* = 4,288 proteins). Due to a few extreme outliers on the order of > 10 standard deviations away from the estimated mean, Normal distributions were fit to the middle 95% of differences, *y_i_*s, for each temperature channel *j*. Using the estimated means and standard deviations, we simulated data to overlay the empirical data, the results of which are shown in Figure 3A (gray). The significant overlap from this analysis demonstrates that a Multivariate Normal model may sufficiently represent the empirical data.

As a result, a vector comprised of the differences for each protein, (*y*_*i*,1_, …, *y*_*i*,10_) for a TMT 10-plex, can be modeled as a Multivariate Normal observation, as shown in Figure 3B. In the unique case of Multivariate Normal data, a correlation of approximately zero between components implies independence. For example, for a given protein, the difference between the control and treatment fraction of non-denatured protein in the first temperature channel, *y*_*i*,1_, is independent that of the second temperature channel, *y*_*i*,2_, as shown in Figure 2C (right). Taken together, a Multivariate Normal model sufficiently represents the null distribution underlying TPP data and bypasses inherent dependencies.

To assess which proteins fall under the alternative hypothesis, that a given protein interacts with a small molecule of interest, the log of the Multivariate Normal *p*-value ranks the proteins from most likely to least likely shifted. This *p*-value may serve as an alternative statistical test. Conceptually, a smaller (more negative) value indicates that a given protein melting curve is more “rare” among the rest of the data. A nifty advantage of this approach allows internal correction and comparisons across all of the data. Because 10-dimensional space is very large, individual *p*-values are also calculated for each *y_i,j_*. Individual *p*-values allow for inspection of the multivariate *p*-value, a quality control checkpoint which would otherwise not be available from a single collapsed statistic, as in the case of PISA^13^. As proteins exhibit a continuous spectrum of curves ranging from unaltered to extremely shifted, a user-defined percentile threshold is recommended to choose the top most likely shifted proteins.

Furthermore, to address noisy protein curves, the variance is calculated for all Multivariate Normal observations, or all *y_i,j_*s for a given protein *i*. The variance peaks near zero and is skewed right, as expected. A warning for high variance is issued if the variance of a Multivariate Normal observation for a given protein is greater than two standard deviations away from the mean variance. While this threshold is qualitative, the warning intends to highlight curves that oscillate between highly positive and negative values.

We have written this analysis in R and as a Shiny App. Notably, the Shiny interface requires no prior knowledge of R or coding and can be used with relative ease. The R code is open-source and available online at https://github.com/alex-bio/TPP, and the Shiny App is available online at https://gygi.med.harvard.edu/software. The Shiny App allows the user to download the result list as an Excel file and has an in-app curve viewer. All of these features make this analysis method accessible to all.

To validate this method, the gold standard panobinostat TPP dataset was examined. Rank-ordering by the log of the Multivariate Normal *p*-value, the previously identified direct and indirect targets of panobinostat^11^ fell within the top 3% most shifted proteins, as shown in Figure 4 (first 7 curves, left to right). We note that HDAC6 and HDAC8, two known panobinostat targets, ranked lower, in the top 17% and 15%, respectively. This indicates that the shifts demonstrated by HDAC6 and HDAC8 are more common across all proteins, which is supported by the slight shift in melting profiles. Furthermore, above the same 3% threshold, many other targets were identified, as shown in Figure 4 (subsequent 9 curves, left to right). Among the other targets are NHLRC3, LAMTOR2, and ASAH1, which are involved in myeloid leukocyte activation and may offer further insight into multiple myeloma therapeutics.

**Figure 4.**
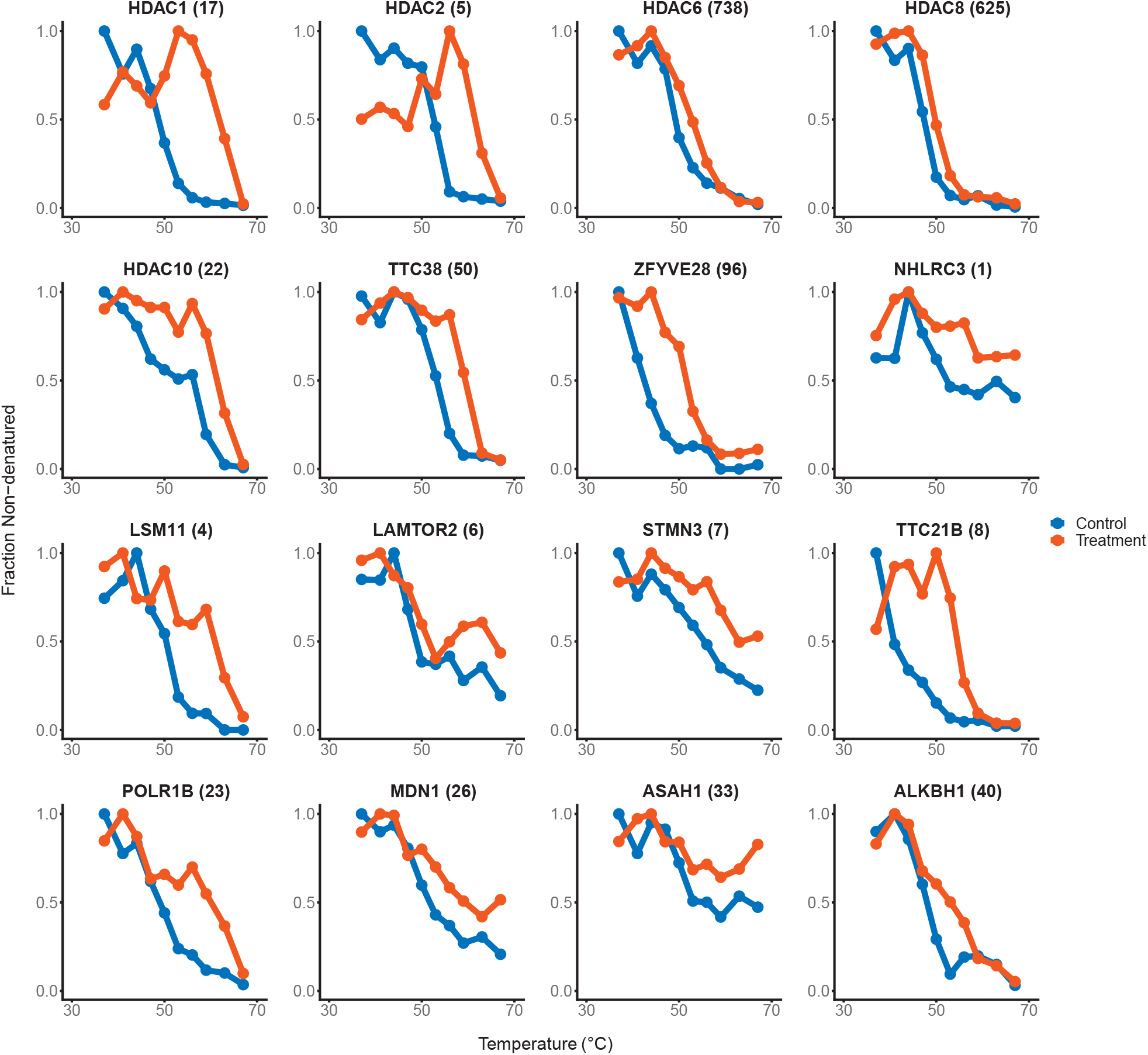
Modeling the panobinostat TPP data^6^ as Multivariate Normal. The Multivariate Normal *p*-value rank, indicating most to least likely shifted, is shown in parentheses after each protein name (*n* = 4,288 proteins). The previously known targets of panobinostat^11^, HDAC1, HDAC2, HDAC10, TTC38, and ZFYVE28, rank in the top 3% of most likely shifted proteins. HDAC6 and HDAC8 ranked lower, in the top 17% and 15% respectively. The melting profiles of identified novel targets, NHLRC3, LSM11, LAMTOR2, STMN3, TTC21B, POLR1B, MDN1, ASAH1, and ALKBH1, are shown. NHLRC3, LAMTOR2, and ASAH1 are involved in myeloid leukocyte activation.

## Conclusion

We present AID, a TPP analysis capable of handling all types of protein melting curves. This method approximates the underlying null distribution, which excludes dependencies within conditions. Moreover, this method compares proteins against the entire dataset rather than pairwise or individually, allowing for a more precise perspective. Applying the analysis to an experiment using panobinostat, a multiple myeloma therapeutic, identified additional protein candidates that may offer insight into malignant mechanisms. This method can be extended to any number of temperature channels, ligands or protein interactors, or general curve data for deeper biological insight.

## Acknowledgements

We would like to thank Jeremy O’Connell and Joao Paulo for considerable mentorship; David Nusinow, and Ed Huttlin for thoughtful feedback and discussion; and funding from the NIH, under award number H667945.

